# RPcontact: Improved prediction of RNA-protein contacts using RNA and protein language models

**DOI:** 10.1101/2025.06.02.657171

**Authors:** Jiuhong Jiang, Xing Zhang, Jian Zhan, Zhichao Miao, Yaoqi Zhou

## Abstract

Determining key contacts between RNA-protein interactions is essential for understanding the molecular mechanisms of numerous biological processes, including transcription, splicing, and translation. However, progress in this area has been impeded by the scarcity of RNA-protein complex structures in the Protein Data Bank (PDB) and the challenges posed by traditional structural determination techniques. Recent computational advancements, including deep learning methods like AlphaFold 3 and RoseTTAFoldNA, have improved contact prediction but are still limited by the availability of homologous sequences and templates. Here, we introduce RPcontact, a novel computational method designed to predict RNA-protein contacts using large language models tailored for RNA (ERNIE-RNA) and proteins (ESM-2). Despite being trained entirely on ribosomal RNA-protein (rRNA-protein) complexes, RPcontact demonstrates robust and generalized performance in predicting contacts for both dimeric and multimeric non-rRNA-protein complexes. The performance of RPcontact on contact predictions significantly improves over the binary contacts inferred from RNA-protein complex structures predicted by AlphaFold 3 and RoseTTAFoldNA, highlighting its potential in RNA-protein complex structure and function prediction.

## Introduction

60 years ago, the heterogeneous nuclear ribonucleoproteins (hnRNPs) research^1^ by James Darnell and colleagues unveiled the intricate functional orchestra of RNA-protein interactions, yet our understanding of molecular mechanisms of RNA-protein binding remains far from complete. It has been shown that non-coding RNAs, at least one order of magnitude more than proteins^2^, can function through interacting with proteins in many processes, ranging from RNA splicing, tRNA synthesis, RNA transcription, and ribozyme activities that are often dependent on protein interactions^3^. The best way to understand how RNA interacts with protein is to solve its complex structure. However, due to physio-chemical properties of RNAs, their structures are more difficult to determine by traditional techniques such as nuclear magnetic resonance^4^, X-ray crystallography^5^ and cryo-electron microscopy^6^, with only 6,029 RNA-protein complex structures, compared to 130,790 protein-protein complex structures available in Protein Data Bank (PDB)^7^ as of September 10, 2024.

Computational methods were developed in genome-wide prediction of RNA binding proteins (RBPs) ^8,9^, binary interactions between RBPs at the whole molecular level^8,10–16^ as well as RNA-binding sites in proteins^17–24^ and protein-binding sites in RNAs^21,25,26^, with the consideration of partner information or RNA preference^27,28^. catRAPID omics v2.0^29^ attempted to predict binding regions by searching for interaction motif pairs in their database. Only a few methods directly predicted RNA-protein contacts at the nucleotide-residue level. lasso^30^ and PRactIP^31^ predicted RNA-protein contacts using machine learning, with the former relying heavily on homologous sequences and the latter on the prediction accuracy of protein/RNA secondary structures. Both methods achieved relatively low accuracy in contact prediction. EV_RNA^32^ predicted RNA-protein contacts from the paired homologous sequences by analyzing the co-evolution between an RNA and a protein, but the quality of the paired homologous sequences has limited the prediction accuracy.

Recently, deep learning methods have brought a revolution in accurately predicting inter/intra-chain protein contacts^33–35^ or RNA contacts^36,37^. These innovations have further extended to the prediction of RNA-protein complex structures, exemplified by the latest developments in AlphaFold 3^38^ and RoseTTAFoldNA^39^. These powerful tools hold great promise for improving sequence-based predictions of RNA-protein contacts at the nucleotide-residue level. However, despite these advancements, the performance of these methods remains limited due to the scarcity of homologous sequences and the limited availability of experimentally determined RNA-protein complex structures in the Protein Data Bank (PDB)^7^.

Recent breakthroughs in large language models, such as RGN2^40^, ESM-2^41^, RNA-FM^42^ and ERNIE-RNA^43^ indicate that these models may inherently capture evolutionary principles within their embeddings, without relying on homologous sequence information. This suggests a novel approach to enhancing the prediction of intermolecular contacts between RNA and proteins. In this study, we introduced a method for predicting RNA-protein intermolecular contacts based on primary sequences, utilizing large language models tailored for RNA (ERNIE-RNA) and proteins (ESM-2). Our algorithm, named RPcontact, has demonstrated robust accuracy and superior performance across various independent test datasets, outperforming existing methods, including AlphaFold 3.

## Results

### RNA-protein contact prediction with RPcontact

To ensure the generalizability of the RPcontact (the schematic framework was shown in **Fig. 1**), we trained RPcontact on 511 ribosomal related RNA-protein (rRNA-protein) complexes only, validated by 49 non-redundant rRNA-protein complexes (VL), and tested by two test sets of non-rRNA-protein complex structures [multimeric (TS1) and dimeric (TS2) complexes] (see Methods). **Fig. 2a** displays receiver Operating Characteristic (ROC) curves for VL, TS1, and TS2 sets, which are nearly overlapping. The respective areas under the ROC curve (auROC) values are 0.85, 0.82, and 0.82. This consistent performance across the rRNA-based validation set and the rRNA-excluded test sets underscores the model’s robustness and its ability to generalize.

**Figure 1.**
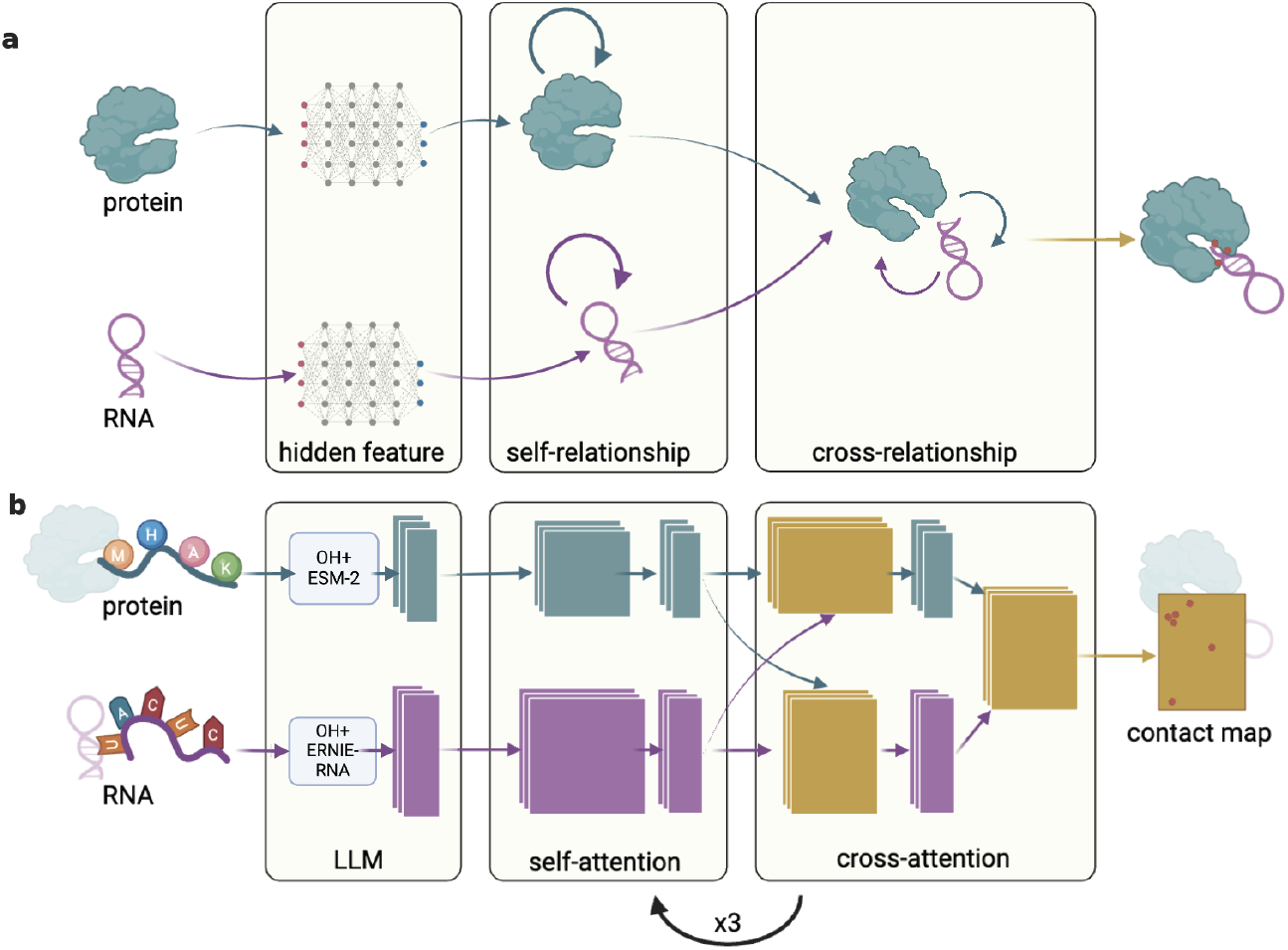
The overall framework of RPcontact. **a.** Conceptual design of RPcontact, where self-relations within a protein or an RNAs are employed for prediction of inter-relationships between RNA and protein. **b.** Overview of RPcontact: An end-to-end prediction of RNA-protein contact map by inputting primary RNA and protein sequences. The program consists of three modules. The first module employed the input primary sequences for extracting embeddings (hidden features) from Large Language Models (LLMs), specifically ERNIE-RNA for RNA sequences and ESM-2 for protein sequences. These embeddings combine one-hot encoding into the second module, which utilizes self-attention to learn the internal relationships within RNA or protein sequences (self-relationship) independently. The third module integrates the information jointly (cross-relationship) through cross-attention. The last two modules are updated three times in series to explore the interactions among all site relationships. Finally, the outputs are concatenated into a matrix representing RNA-protein contacts, determined by applying a threshold.

**Figure 2.**
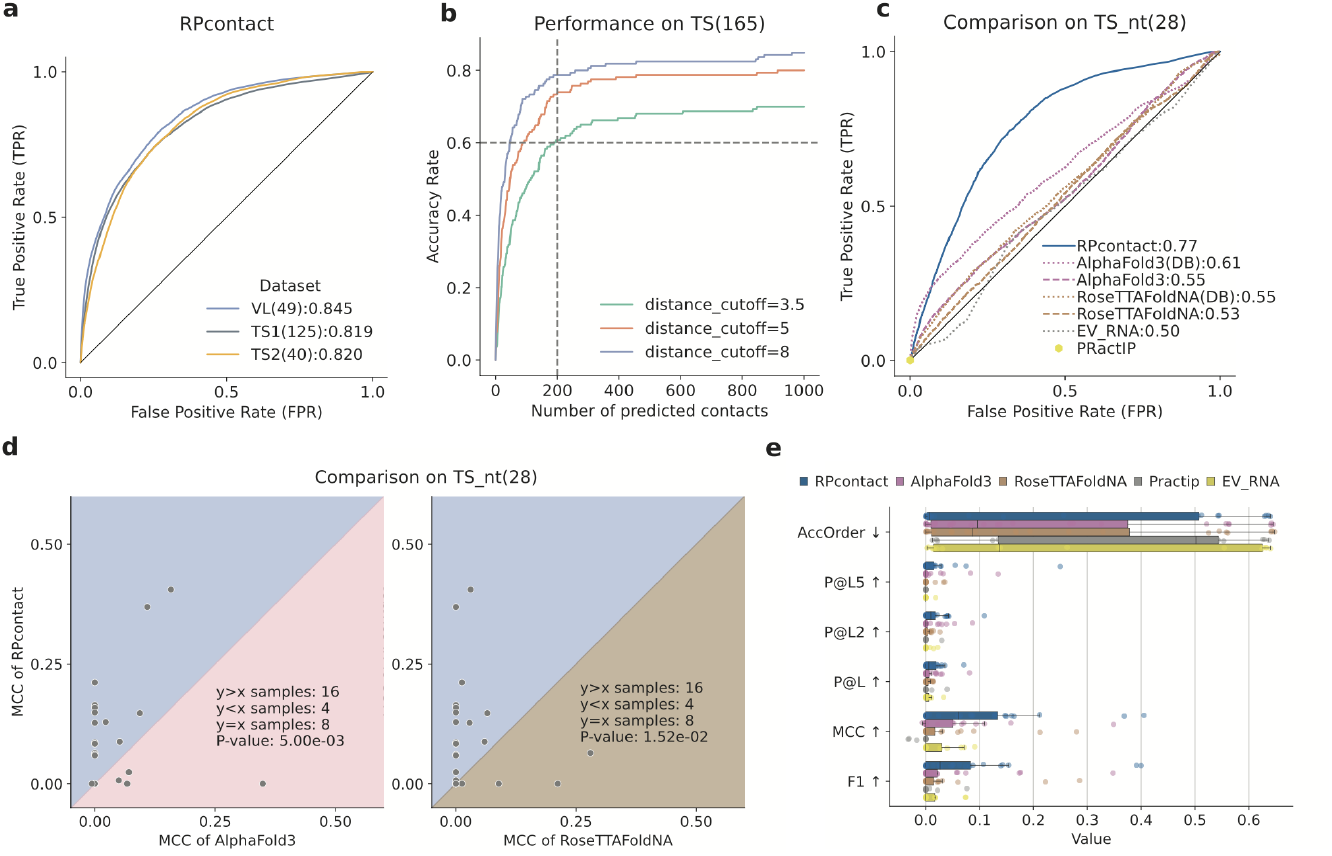
Robustness of RPcontact performance across multiple test sets along with comparison to other methods. **a.** Similar receiver operating characteristic curves (ROCs and area under ROCs, auROCs) on the validation set of rRNA-protein complexes and two independent test sets of multimeric (TS1) and dimeric (TS2) non-rRNA-protein complexes given by RPcontact indicates robust and generalizable performance. **b.** RPcontact performance in terms of accuracy rate in three distance cutoffs as a function of number of predicted contacts for the combined test set TS (TS1+TS2). **c.** The ROC curve given by RPcontact is compared to several methods on TS_nt (28 non-redundant targets against the training set of AlphaFold 3) with 26% higher auROC than the second best AlphaFold 3(DB). **d.** Performance comparison on individual RNA-protein contact maps according to AccOrder (the smaller the better) between RPcontact and AlphaFold 3 and between RPcontact and RoseTTAFoldNA on TS_nt. **e.** Performance comparison according to AccOrder, MCC, precision under different cutoffs (top L/5, L/2, L predictions), and F1-score, L is the sum length of rna and protein sequence. The point plot illustrates the central tendency and variability of different metrics across various values, categorized by a metrics. Each point represents the average mean for a specific metric value, with error bars indicating the uncertainty of the estimate. The error bars represent the confidence intervals of the means, while the points in the middle of the error bars indicate the mean values, reflecting the performance and uncertainty across different methods.

To further evaluate the model’s accuracy, we analyzed the prediction results using different distance cutoffs. **Fig. 2b** compares the accuracy rates of targets with correctly predicted contacts evaluated with different distance cutoffs (3.5Å, 5Å, 8Å) as a function of the number of predicted contacts. Although the model was trained with 5 Å cutoff, the result indicates that the accuracy rate is higher with a more relaxed definition of contact, as expected. For example, 61% of the complexes in the test set (TS) would give at least one true positive contact for a distance cutoff 3.5 Å, 74% for 5 Å cutoff and 79% for 8 Å cutoff at top 200 predictions. This indicates that RPcontact can reduce the search space for true positive contacts. That is, even if a contact may be a false positive at the 5Å cutoff, it could be a true contact at a longer distance cutoff.

RPcontact was benchmarked against other methods on TS_nt (28 non-redundant targets against the training set of AlphaFold 3 and other methods), including the structure prediction method AlphaFold 3 and RoseTTAFoldNA with or without employing the database for homologous sequences and templates (AlphaFold 3 and RoseTTAFoldNA versus AlphaFold 3(DB) and RoseTTAFoldNA(DB), respectively), the coevolution-based method EV_RNA, and the contact predictor PRactIP. As shown in **Fig. 2c,** EV_RNA and RoseTTAFoldNA both yielded a performance equal to or slightly better than random with auROC at 0.50 and 0.53. Only one true positive contact was predicted by PRactIP in these 28 targets, giving a true positive rate near to 0. There was a large drop in performance for AlphaFold 3 and RoseTTAFoldNA without homologous sequences or templates (auROC reduces from 0.61 to 0.55 for AlphaFold 3 and 0.55 to 0.53 for RoseTTAFoldNA). Notably, RPcontact does not require the extensive computational resources typically associated with database-dependent methods, offering a significant advantage in scalability and efficiency. RPcontact leads with an AUC of 0.77, even surpassing the methods with homologous sequences and templates like AlphaFold 3(DB) and RoseTTAFoldNA(DB). The MCC comparisons in **Fig. 2d** highlight RPcontact’s superior performance, with a P-value of 5.00e-03 against AlphaFold3 and 1.52e-02 against RoseTTAFoldNA, indicating statistically significant improvements (P-value < 0.05). RPcontact enhanced the prediction for the targets that AlphaFold3 or RoseTTAFoldNA predicted with near-zero MCC scores. The head-by-head comparison to AlphaFold3(DB) and RoseTTAFoldNA(DB) is shown in **supplementary Fig. S1**. RPcontact still significantly outperforms AlphaFold3(DB) and enhances the targets when AlphaFold3(DB) loses its predictability.

Additional metrics, including AccOrder, P@L5, P@L2, P@L, MCC, and F1 score, were benchmarked on TS_nt for the five prediction algorithms in **Fig. 2e**. RPcontact consistently yields the lowest AccOrder and the highest P@L5, P@L2, P@L, MCC, and F1 scores in terms of the median value. Furthermore, the head-by-head comparison for all these metrics, provided in **supplementary Fig. S2**, yielded the same conclusion.

### Large language model facilitates contact prediction in RPcontact

To examine the factors that might have contributed to the performance of RPcontact, we also trained a series of model variants in different groups.

The “Baseline” group in **Table 1** (and **Supplementary Fig S3a**) compares the performance of RPcontact according to auROC in the absence of large language models (LLMs) (i.e., one-hot-encoding only (OH)), as well as in cases where the LLM is partially missing (protein (P_Emb) only or RNA embedding (R_Emb) only). All model variants were trained in the absence of data augmentation at the distance cutoff of 5Å. RPcontact with both LLMs (OH+RP_Emb) provides a substantial improvement over OH, OH + R_Emb, OH+ P+Emb across validation and two test sets. In fact, without data augmentation, RPcontact with both LLMs is the only method that consistently achieves an auROC > 0.79 across all three datasets. **Supplementary Fig S3b** provides a further comparison under the condition of employing data augmentation, contrasting the use of one-hot-encoding alone (OH) with RPcontact combined with both LLMs. The significant improvement also shows the power of employing LLMs.

**Table 1.** Performance on the model variants. Comparison of auROC when training on ribosome data, validating on ribosome data (VL), and testing on other data (TS1, TS2).

| Group | Setting | Variate | Name | VL(49) | TS1(125) | TS2(40) |
| --- | --- | --- | --- | --- | --- | --- |
| Baseline | w./o.<br>augm.<br>Distance<br>cutoff=5 Å | w./o. LLM | OH | 0.776 | 0.753 | 0.693 |
|  |  | RNA LLM | OH+R_Emb | 0.777 | 0.729 | 0.707 |
|  |  | Protein LLM | OH+P_Emb | 0.826 | 0.779 | 0.756 |
|  |  | w./ LLM | OH+RP_Emb | 0.841 | 0.801 | 0.793 |
| Distance cutoff | w./ augm.<br>w./ LLM | 3.5 Å | cutoff=3.5 | 0.830 | 0.810 | 0.806 |
|  |  | 8 Å | cutoff=8 | 0.761 | 0.740 | 0.737 |
|  |  | 5 Å | RPcontact | **0.845** | **0.819** | **0.820** |
| Data augm. | w./ LLM<br>Distance<br>cutoff=5 Å | w./o. augm. | TR(511) | 0.841 | 0.801 | 0.793 |
|  |  | w./ augm. | RPcontact | **0.845** | **0.819** | **0.820** |
| Language model | w./ augm.<br>w./ LLM<br>Distance<br>cutoff=5 Å | RNA-FM | OH+RP_Emb | 0.838 | 0.782 | 0.764 |
|  |  | ERNIE-RNA | RPcontact | **0.845** | **0.819** | **0.820** |

Note:
1. The number in parentheses next to a dataset represents the number of instances in that dataset. 2. The "improve" column is the improvement percentage of auROC on VL by comparison with the baseline variant. 3. "+" signifies concatenating different features before inputting them into the model. 4. R\_Emb. refers to each RNA sequence being encoded using the embedding extracted from the RNA-FM model or the ERNIE-RNA model. 5. P\_Emb. refers to each protein sequence being encoded using the embedding extracted from the ESM-2 model. 6. w. augm. indicates that data augmentation is utilized, and 3,593 instances are employed for training. Otherwise, w./o. augm. is set to FALSE, and only 511 full-length instances are used. 7. The dataset of the training set is ribosome data, whereas the remaining datasets are not. 8. w./, with, w./o, without 9. LLM. large language model.

The “Distance cutoff” group in **Table 1** (and **Supplementary Fig S3c**) explores the influence of varying distance cutoffs on the generation of training labels. Specifically, an increase in the distance cutoff from 3.5 Å to 5 Å results in a modest improvement of approximately 2%. However, extending the cutoff from 5 Å to 8 Å leads to a significant decline of around 11%. Additionally, we assessed the impact of data augmentation (as described in the “Data augm.” group in **Table 1** and **Supplementary Fig S3d**). While data augmentation only provided a minor improvement on the VL set (0.5% increase in auROC), it yielded a more pronounced improvement of 2% on the TS1 and 3% on TS2. This suggests that data augmentation is effective in generalizing the rules learned from rRNA-protein complexes to non-rRNA-protein complexes.

The “Language model” group in **Table 1** examines the comparative effects of two RNA language models, namely RNA-FM and ERNIE-RNA. The primary distinction between the two LMs is that ERNIE-RNA takes into account the potential of base pairs, particularly the lower energy of AU pairs compared to CG pairs, and incorporates that pairing information into its attention mechanism. Here, the results demonstrate that ERNIE-RNA outperformed RNA-FM across all three datasets (VL, TS1, and TS2). Nevertheless, employing only the RNA language model, without incorporating a protein-language model, failed to produce a significant improvement in prediction performance. This contrasts with the use of a protein language model alone, as shown in the “Baseline” group in **Table 1** and **Supplementary Fig S3a**, which did result in notable improvements.

### Predictive in non-ribosomal RNA-protein complexes

RPcontact, which was trained on ribosomal RNA-protein complexes, has demonstrated its efficacy in predicting contacts in non-ribosomal RNA-protein complexes. As evidence, **Fig. 3** compares the top 50 predicted contacts of three RNA-protein dimeric complexes against a gold standard defined by distance cutoffs at 3.5 Å, 5 Å, and 8 Å. These complexes include 3KTW_D_B^44^ (SRP RNA-SRP19 binary complexes, **Figs. 3a-c**), 8CSZ_C_D^45^ (ωRNA-IscB binary complexes, **Figs. 3d-f**), and 2ZZN_D_B^46^ (tRNACys-aTrm5, **Figs. 3g-h**). RPcontact successfully predicted more than three key contacts (at a distance cutoff of 3.5 Å) for each non-ribosomal RNA-protein complex (**Figs. 3a, 3d, and 3g**). The surface shape of the accurately predicted contacts provided an interface that can guide the docking process of the two components (**Figs. 3b, 3e, and 3h**). Additionally, spheres of the backbone atoms further illustrate the spatial relationships between RNA and protein sites during contact formation **(Figs. 3c, 3f, and 3i**).

**Figure 3.**
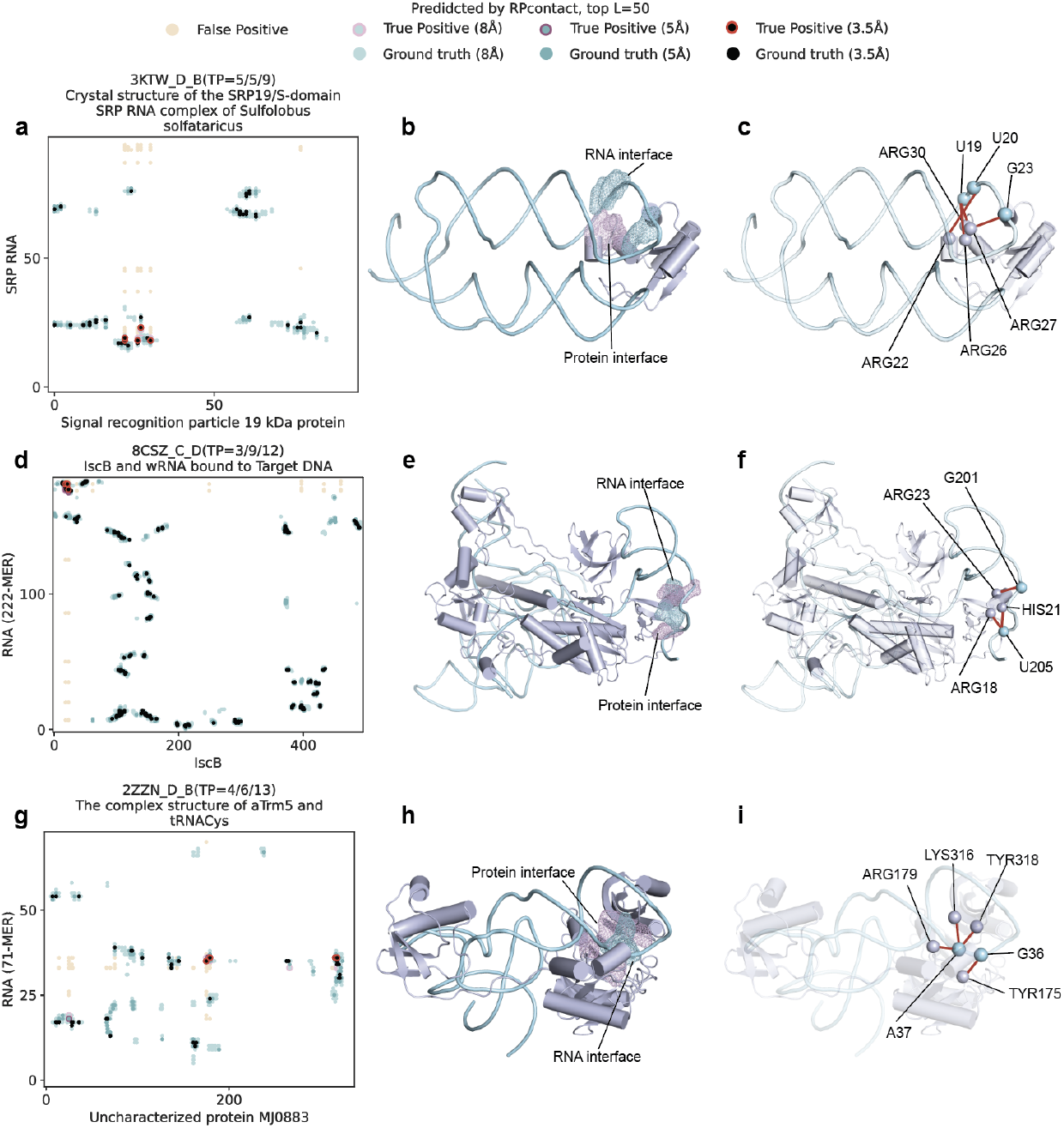
Case studies of non-ribosomal RNA-protein complexes. This figure presents an in-depth examination of three non-ribosomal RNA-protein complexes, arrayed in rows. The complexes include the SRP RNA-SRP19 dimer (panels **a-c**), the ωRNA-IscB dimer (panels **d-f**), and the tRNACys-aTrm5 dimer (panels **g-i**). The rectangles in panels (**a,d,g**) represent contact maps, which are R x P matrices illustrating the interactions between an RNA of R nucleotides (on the y-axis) and a protein of P residues (on the x-axis). The dark, cyan, and light cyan dots within these rectangles signify actual contacts at varying distance thresholds (3.5 Å, 5 Å, and 8 Å), as validated by the Protein Data Bank. Dots with a circled perimeter indicate the contacts accurately predicted by the RPcontact algorithm (True Positives, TP, top 50 as predicted contacts). The wheat-colored dots represent incorrectly predicted contacts (False Positives, FP) across all distance thresholds. Notably, red circles or lines are used to emphasize the key contacts (at a 3.5 Å distance cutoff) that were correctly predicted. The 3D structures of the RNA-protein dimers are visualized in panels (**b,e,h**), with the RNA depicted in cyan and the protein in purple. Interfaces of the truly predicted contacts are accentuated in a surface representation, and the backbone atoms (P for nucleotides and Cα for amino acids) are annotated on spheres in panels (**c,f,i**). Each sub-figure title includes details derived from the PDB, such as “3KTW_D_B”, indicating the PDB identifier “3KTW”, with the RNA chain labeled “D” and the protein chain labeled “B”. The notation TP=5/5/9 signifies the number of true positives detected at distance cutoffs of 3.5 Å, 5 Å, and 8 Å, respectively.

### Identification of functional riboregulation sites

RPcontact has successfully predicted the functional riboregulator sites, as demonstrated by its utility in analyzing tRNA-protein complexes as illustrated in **Fig. 3g**. The interaction between tRNACys and the archaeal Trm5 enzyme (aTrm5), which involves an intricate network of binding sites on both the RNA and protein components. The predicted RNA binding sites on the protein are primarily centered around residues 35, 90, 180 and 320, while the predicted RNA binding sites are positioned near nucleotides 20 and 35. RPcontact effectively pinpoints these binding regions on both the RNA and protein sides. Additionally, some false positives are in close vicinity to true positives, indicating their potential role in steering the complexes towards the correct orientation for docking simulations. Within the top 50 predictions, four true positive (TP) contacts were obtained at a distance cutoff of 3.5 Å: A38: ARG181, G37: TYR177, A38: LYS318, and A38: TYR320 (nucleotide position: residue position). Although these contacts are far apart in sequence, such as ARG181 and TYR320, they are spatially adjacent in the 3D structure, as depicted in **Figs. 3h-i**. The protein binding sites associated with these contacts are located in the anticodon stem (P3) of the tRNACys cloverleaf structure, as shown in **Fig. 4**. Notably, nucleotides A38 and G37 are positioned in close proximity to the anticodon (positions 34-36 in the RNA), underscoring RPcontact’s ability in predicting the functional riboregulator sites that are vital for enzyme modification during the functional maturation of this tRNA.

**Figure 4.**
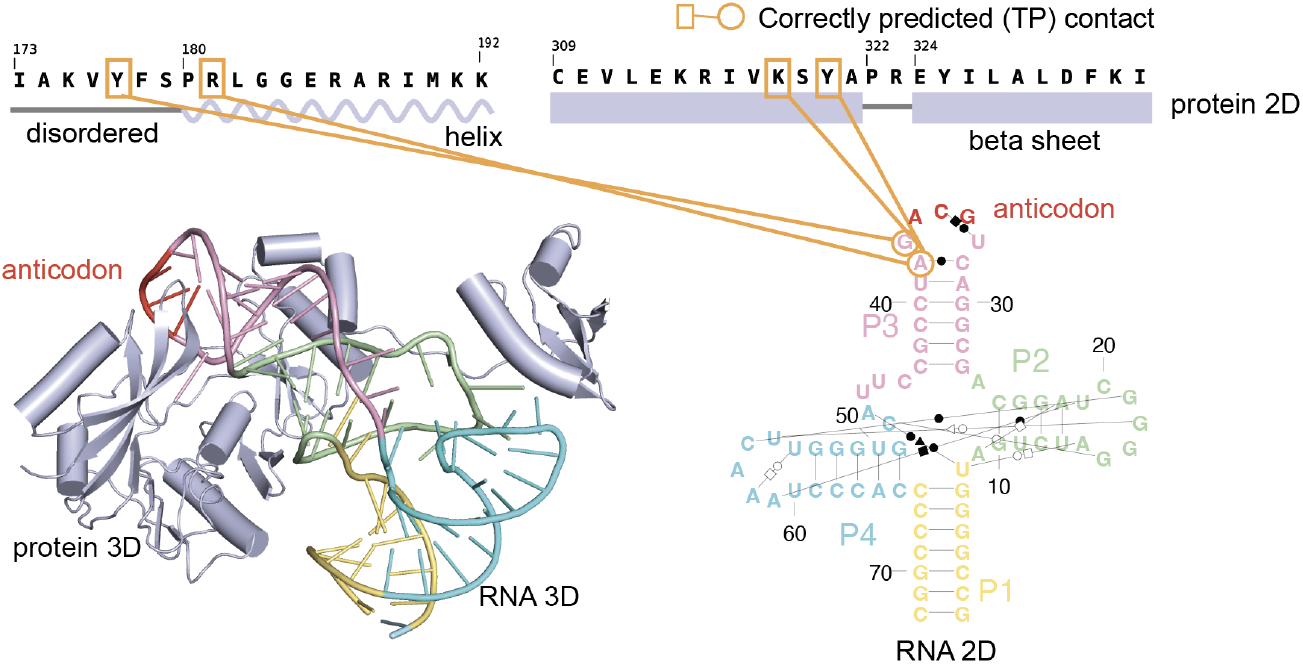
RPcontact identified the key contacts in the complex structure. The overall 3D structure of aTrm5 and tRNACys (2ZZN_D_B) was shown by Pymol, with purple parts representing protein while the other colorful parts represent RNA. The 2D structure of the RNA was presented in a cloverleaf shape with corresponded colors. The anticodon is highlighted in red while the four stem regions are colored differently. The orange circles connected by line represent TP contacts. The TPs are associated binding sites located in the disorder region, helix region and beta sheet region of protein and the P3 region of RNA. TP. true positive.

### Identifying the contacts in multimeric structures

RPcontact predicts contacts within dimeric complexes (a single RNA and protein within a PDB assembly unit) even under the influence of unknown third partners. Multimeric structures provide those case studies. In the prediction process, RPcontact accepts one RNA sequence and one protein sequence as inputs. Therefore, the effect of a third unknown partner(s) increases the difficulty of prediction. **Fig. 5** illustrates the successful prediction of interactions between u1 snRNA (chain P) and sn ribonucleoprotein-associated proteins B (chain b), Sm D1, and u1 sn ribonucleoprotein C (chain R). The 3D structures and ground truth contacts are derived from the PDB (5ZWN) ^47^. **Fig. 5a** presents both the assembly and monomer global views of the biomolecule structures, while the local spatial interactions are shown in **Figs. 5b-d**. In which the backbone atoms of the nucleotides and amino acids are highlighted in a sphere to visually represent their spatial positional relationships. Furthermore, **Figs. 5e-g** shows the correctly predicted contacts at different distance cutoffs (3.5 Å, 5 Å, and 8 Å). For each dimeric RNA-protein complex **(Figs. 5e-g**), RPcontact accurately identifies two contacts when the distance cutoff is 3.5 Å, confirming the method’s generalizability beyond dimeric complexes. This robust performance in predicting complex interactions, even with incomplete information about interacting partners, underscores the model’s strength in versatility and its potential for broader application in biomolecular studies.

**Figure 5.**
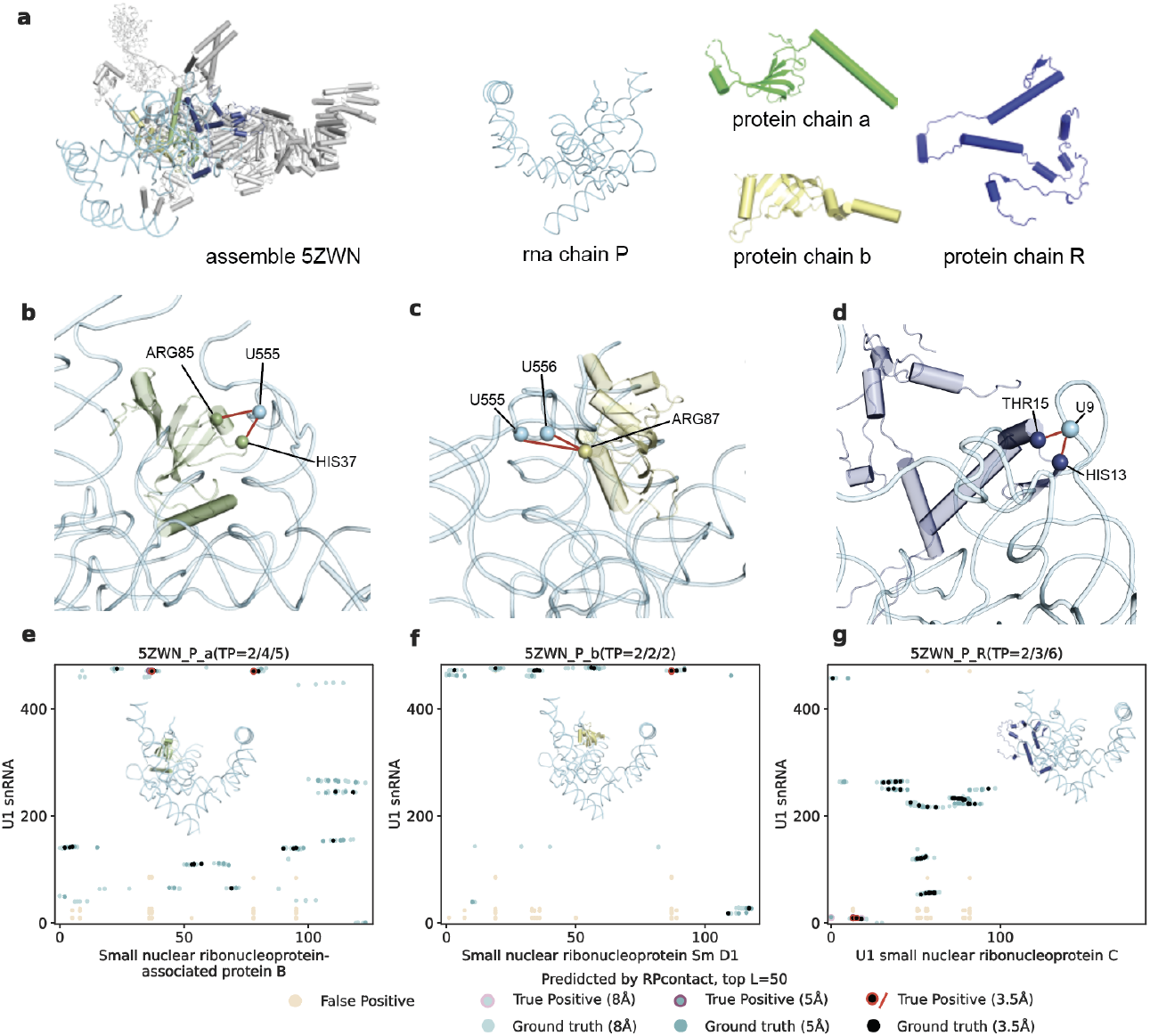
Example of multimer RNA-protein complexes. **a.** Cartoon view of structures. The structures (PDB ID: 5ZWN) are presented in a functional unit (or assembly). The gray regions represent molecules in the assembly that are not investigated in this example. The cyan molecule represents the target RNA (Chain P, small nuclear ribonucleoprotein G), while the other three colored molecules are proteins: Chain a (small nuclear ribonucleoprotein-associated protein B) in green, Chain b (small nuclear ribonucleoprotein Sm D1) in yellow, and Chain R (U1 small nuclear ribonucleoprotein C) in blue. **b-d.** Local structures in surface shape surrounding TP contacts. TP contacts are using red lines to link the two backbone atoms (P for nucleotides and Cα for amino acids), which were depicted in a sphere shape. **e-g.** Key contacts prediction in the contact map. It represents contact maps, which are R x P matrices illustrating the interactions between an RNA of R nucleotides (on the y-axis) and a protein of P residues (on the x-axis). The dark, cyan, and light cyan dots within these rectangles signify actual contacts at varying distance thresholds (3.5 Å, 5 Å, and 8 Å), as validated by the Protein Data Bank. Dots with a circled perimeter indicate the contacts accurately predicted by the RPcontact algorithm (True Positives, TP, top 50 as predicted contacts). The wheat-colored dots represent incorrectly predicted contacts (False Positives, FP) across all distance thresholds. Notably, red circles or lines are used to emphasize the key contacts (at a 3.5 Å distance cutoff) that were correctly predicted. In the space of the contact map, there is the dimer view of the 3D structures for the examined RNA-protein complexes. Each sub-figure title includes details derived from the PDB, such as “5ZWN_P_a”, indicating the PDB identifier “5ZWN”, with the RNA chain labeled “p” and the protein chain labeled “a”. The notation TP=2/4/5 signifies the number of true positives detected at distance cutoffs of 3.5 Å, 5 Å, and 8 Å, respectively.

## Discussion

Ribosomes consist of tens of proteins and several large RNA structures, making them complex molecular machines. Given this complexity, the RNA-protein interactions derived from ribosomes across different species are likely to encompass fundamental principles governing RNA-protein binding contacts. RPcontact supports this notion by training on ribosomal RNA-protein contacts and subsequently testing on non-ribosomal contacts. The model’s success in identifying key positive contacts indicates that the underlying RNA-protein interaction mechanisms are conserved across diverse complexes. Furthermore, our findings show that incorporating repeat local contact maps significantly enhances the model’s ability to generalize to non-ribosomal complexes.

The integration of large language models (LLMs) into the prediction pipeline offers a significant advantage by encapsulating the evolutionary information typically derived from multiple sequence alignments (MSAs). This evolutionary context is crucial for downstream tasks such as contact prediction, functional inference, and structure prediction. Consequently, LLMs can effectively substitute the need for MSAs of homologous sequences, thereby streamlining prediction tasks that rely heavily on evolutionary information. The state-of-the-art performance of RPcontact demonstrates that incorporating both RNA and protein language models supersedes the use of a language model on one side (OH+R_emb, OH+P_emb) as well as a one-hot encoding model (OH) in the absence of language model embeddings. This indicates that language models may help in predicting contacts from primary sequences alone, particularly orphan proteins or RNAs lacking homologous sequences.

RNA language model alone in the absence of a protein language model did not bring a significant improvement for RNA-protein contact map prediction. This suggests the difficulty in extracting binding sites information from the embedding without corresponding embedding from its partner, but the opposite is not true (protein language model alone can help). Such difficulty may be because the RNA sequence is not as conserved as a protein due to the small number of letter codes (only 4 versus 20) and correlated mutations. RNA sequences can tolerate certain base pair substitutions, such as AU pairs interchanging with UA pairs, without disrupting the overall RNA structure. Thus, an improved RNA language model with more accurate capture of the substitutions and base pairs in 2D structure will likely further improve prediction in RNA-protein interaction contacts.

Comprehensive benchmarking underscores RPcontact’s superior performance and efficiency, making it a highly valuable tool for RNA-protein structure prediction, particularly in scenarios where computational resources are limited. RPcontact not only demonstrates an enhanced ability to capture the interactions between nucleotide bases and amino acid residues but also plays a pivotal role in complex docking stimulation. A detailed comparison of the ground truth and predicted contact maps for nine complexes in the TS_nt dataset is provided in **Supplementary Fig. S4** and **Fig. S5**, showcasing the predictions made by RPcontact and AlphaFold 3, respectively. RPcontact successfully identified contacts with a distance cutoff of 3.5 Å for all 9 complexes, while AlphaFold 3 lost predictability in 6 out of 9 complexes (no true positive contacts identified). The detection of contacts is particularly significant as it effectively restrains most conformations for a given dimer, thereby reducing the conformational space that needs to be explored during docking simulations. By providing more precise and reliable contact information, RPcontact enables efficient reduction of the conformational space for complexes with low demands for computational resources.

Thus, RPcontact is able to provide a quick, approximate prediction of the RNA-protein contact maps from large-scale datasets, thereby addressing the gap in our understanding of the functions of sequences within the vast genome. For example, applying RPcontact to the eCLIP^48^ dataset, we could fully leverage the experimentally validated interactions to annotate RNA-protein binding interactions at the site-specific level. This approach should allow us to uncover the complex interdependence between the cellular transcriptome and proteome in unprecedented detail. Rather than relying on broad binding regions, RPcontact could pinpoint the interaction sites between RNA and protein sequences or structural elements. This enhanced resolution would provide valuable mechanistic insights into the molecular basis of RNA-protein complex recognition and regulation. Identifying the specific interaction sites can shed new light on how RBPs recognize and bind to their target RNAs, and how this binding contributes to diverse cellular processes. Bridging the gap between genome sequence information and structural determinants of RBP-RNA complex formation remains an important challenge in the field. Working in this area is in progress.

RPcontact prediction may also be useful for mRNA sequence design. RBPs are known as the major regulators of both coding and non-coding transcripts^49^, with examples including translation initiation, elongation and termination factors, as well as proteins involved in RNA degradation processes. When designing mRNA sequences for various applications such as therapeutics, it is crucial to identify and eliminate the specific interaction sites between the mRNA and regulatory proteins that could lead to undesirable binding and degradation. By pinpointing the possible contact points between the mRNA and factors like RNA decay machinery, one can rationally engineer the mRNA sequence to avoid these detrimental interactions, thereby optimizing mRNA stability and ensuring effective translation. This targeted approach to modulating RBP-mRNA contacts may provide a powerful strategy for fine-tuning mRNA design and function, in contrast to relying on broader binding regions. Developing accurate computational tools to map these critical RBP-mRNA interaction interfaces remains an important challenge in realizing the full potential of engineered mRNA technologies.

RPcontact predicts RNA-protein contacts directly from the primary sequence, independently of background sequences or structural databases, which are traditionally relied upon by tools like AlphaFold 3 and RoseTTAFoldNA. Thus, this method not only reduces the need for additional database storage and the time-consuming process of searching for similar sequences (homologues) or structures (templates) but also offers significant “lightweight” advantages. The lightweight nature of RPcontact inference simplifies the prediction process, reduces computational resource consumption, and enhances the speed and efficiency of predictions. Furthermore, its ease of use allows researchers to quickly adopt this method for their studies.

## Methods

### Definition of the RNA-protein contact map

RNA-protein contacts can be defined according to experimentally determined complex structure^50^. A contact is formed between residue *i* of a protein and nucleotide *j* of an RNA if any atom in residue *i* is within a cutoff distance (e.g., 5 Å^51,52^, 3.5 Å^53^, 8 Å^32^) of any atom in nucleotide *j*. Here, 5 Å represents possible pi-pi, anion-pi, and cation-pi interactions. 3.5 Å is related to the hydrogen bond distance, while 8 Å can be a potential solvent-mediated interaction distance^54,55^. Thus, the contact map between an RNA of *R* nucleotides and a protein of *P* residues is defined as a binary matrix of *R x P*, in which 1 means a contact and 0 otherwise. The ground truth of all contact maps is obtained from experimentally determined structures from the Protein Data Bank^7^. Here, we employed 5 Å as the cutoff for training and evaluating RPcontact.

### Training, validation, and test datasets

A dataset of RNA-protein complex structures with resolutions of 3.5 Å or better was downloaded from the PDB, including both X-ray Diffraction and Electron Microscopy techniques. It yielded a total of 4,399 assemblies (retrieved before May 10, 2023). This dataset was curated to include only those complexes where both RNA and protein chains were present within the functional assembly, excluding RNA sequences outside the 32 to 1000-nucleotide range. We also removed those containing fewer than four types of nucleotides to avoid possible biases because RNAs containing fewer than four types of nucleotides typically consist of simple sequence repetitions and interact with similar binding sites on multiple identical proteins.

To prevent data leakage between training/validation and testing sets, we removed redundancy in the dataset from both RNA and protein components of the complexes. During the redundancy removal process, complexes with fewer numbers of contacts were removed. From the protein side, sequence redundancy was reduced using psi-cd-hit^56^ with a 30% identity cutoff, utilizing BLAST^57^ for enhanced alignment with the command “cd-hit-v4.8.1-2019-0228/psi-cd-hit/psi-cd-hit.pl -i protein.fasta -o protein_0.3 -c 0.3 -P blast/2.10.0/bin/”. This led to 725 RNA-protein complexes. From the RNA side, molecules whose names include the keywords “ribosome” or “ribosomal” were assigned to the training and validation sets. These sets comprise a total of 560 ribosomal-related complexes (rRNA-protein complexes). They were randomly divided into TR (511) and VL (49) sets. The rest 165 non-rRNA-protein complexes, named TS (the length distribution is shown in **Supplementary Fig. S6**), were further divided into multimeric RNA-protein complexes test set (TS1, 125) and dimeric RNA-protein complexes (one RNA and one protein in a complex, TS2, 40). To ensure a fair comparison with AlphaFold 3, we curated a dataset by initially selecting 11 targets from TS (165), excluding any proteins with homology to those in AlphaFold 3’s training set. Recognizing the limitations of a small dataset, we expanded our collection by incorporating newly deposited structures ranging from May 11, 2023 to November 24, 2024. Utilizing BLAST, we excluded homologous proteins against the protein released before May 11, 2023. This process led to a final dataset comprising 17 targets. Combined with the initial 11 targets from TS, we established a non-redundant dataset (TS_nt, 28 targets).

To relieve the imbalance (shown in **Table S1**) between positive (contacts) and negative (non-contacts) labels during training, we applied a data augmentation technique that repeated the dense local contact map of the targets in TR. We employed random-sized windows across local contact maps. These windows moved in random-sized steps, as shown in **Supplementary Fig. S7**. The window size, map height, and step size were all randomly chosen between 32 and 50. We only adopted local contact maps with more than three contacts.

### The algorithm design of RPcontact

RPcontact employs large language models for the prediction of RNA-protein contacts from their primary sequences. It contains three modules as shown in **Fig. 1**.

The first module takes the sequences as input and employs large language models for producing sequence embeddings. Specifically, we utilize ERNIE-RNA^43^, a model trained on non-coding RNAs from RNA Central, which generates 768-dimensional embeddings for each nucleotide in an RNA sequence to capture functional and evolutionary features. These embeddings, combined with 4-dimensional one-hot encoded nucleotide type information, serve as inputs to our neural network. For protein sequences, we employed ESM-2^41^, a transformer protein language model with up to 15 billion parameters (esm2_t48_15B_UR50D), to produce a 5120-dimensional embedding for each residue. The 5120-dimensional embeddings, together with the 20-dimensional one-hot encoded residue type information, are integrated into our neural network to enhance its predictive capabilities.

The second module, then, captures intra-molecule (RNA or protein) site-to-site relationship with a neural network with a self-attention mechanism, because nucleotide-residue binding is highly related to their respective intra-molecular interactions. In this module, high-dimensional inputs of RNA and protein are condensed into 48 dimensions by a linear layer at the beginning of the second module.

The third module builds on the knowledge of self-relation to infer cross-relationships using a neural network with a cross-attention mechanism. This module amplifies the 48-dimensional vectors from the second module to 512 dimensions, which are then re-condensed to 48 dimensions for reinjection back to the second module. This process iterates three times to comprehensively refine and exchange the inter- and intramolecular relationships between RNA and protein sites, ultimately distilling the outputs into a more concise representation. **Supplementary Fig. S8** shows more details about the neural network in RPcontact.

To limit possible false positives in the outputs of the model, we set a maximum of 24 protein residues per nucleotide and 12 RNA nucleotides per residue. This constraint allows the method to focus on the most significant contacts. Additionally, we normalized the scores of the remaining contacts using max-min normalization. Hence, RPcontact gives a score between 0 to 1 for each residue-nucleotide pair, representing the possibility of the contact forming.

RPcontact was implemented in PyTorch and trained over a total of 38 epochs on an X86 64-bit 2.0GHz CPU with 512GB of memory. In total, the deep neural network in RPcontact includes 696,433 trainable parameters. We utilized the Root Mean Squared Propagation (RMSProp) optimizer ^57,58^ during the training process. The batch size and learning rate were set to 32 and 0.0001, respectively. The model utilized a cosine annealing schedule to progressively reduce the learning rate, with regular restarts intended to prevent it from getting trapped in a local minimum. Training was halted when the binary cross-entropy loss between predicted and actual labels on the validation set did not increase for 10 consecutive epochs.

### Metrics to evaluate the prediction accuracy of RNA-protein contacts

We measure the method’s performance according to the area under the Receiver Operating Characteristic (ROC) curve (auROC), which is a graphical representation of the True Positive Rate (TPR) against the False Positive Rate (FPR) [**eq. (1-2)]** at various threshold settings as below:

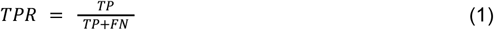

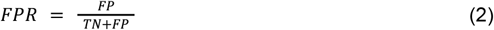

where TP, FN, FP, TN are the number of true positive contacts, false negative contacts, false positive contacts and true negative contacts, respectively. The auROC value, ranging from 0 to 1, quantifies the model’s ability to distinguish between positive and negative contacts. A higher auROC value indicates a better model performance, meaning the model is more effective at generating a higher TPR with a lower FPR. An auROC value of 0.5 indicates a random predictor, while values greater than 0.5 suggest an effective predictor. The auROC is a valuable metric for evaluating the performance of universal binary classification models, particularly in imbalanced datasets.

We also measure the performance by Precision [**eq. (3)**], F1-score [**eq. (4**)], and Matthews Correlation Coefficient (MCC) [**eq. (5**)], as below:

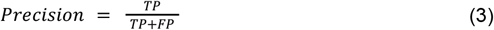

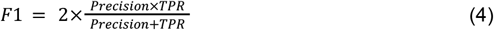

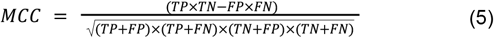

MCC, in particular, is robust against class distribution imbalance, providing a comprehensive evaluation of model accuracy. For precision, we further defined P@L5, P@L2, P@L for evaluating the precision at top L/2, top L/5, top L predictions as has been done previously^59^.

In addition, we employed the accuracy rate, the ratio of correctly predicted complexes in one data set [**eq. (6)]**^**60,61**^

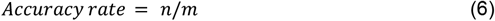

where *n* is the number of targets with at least one true contact among the predicted contacts, and *m* is the number of all targets in one dataset. In this study, one target is one RNA-protein complex. We further defined accuracy order ^60 61^ as shown in **eq. (7)**:

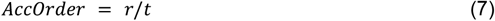

where *r* is the rank of the first positive prediction, and *t* is the total number of base-residue pairing combinations within an RNA-protein complex. A lower AccOrder value signifies a more precise prediction.

The statistical significance (P-value, P-value <0.05 is defined as significant) was assessed using the Wilcoxon signed-rank test, with the “less” option applied for the AccOrder (the smaller, the better) metric while the “greater” option was used for other metrics (the bigger the better) to establish if there was a significant performance improvement.

### Implementation of other methods for comparison

For AlphaFold 3, we submitted sequence pairs (one RNA and one protein sequence) to the server of AlphaFold 3 at https://golgi.sandbox.google.com/. The server returns the predicted 3D structures of RNA-protein complexes. We also downloaded the source code from https://github.com/google-deepmind/alphafold3 and installed AlphaFold 3 locally to obtain the database-free predictions. For RoseTTAFoldNA, the method was installed locally for predicting 3D structures of RNA-protein complexes by downloading from https://github.com/uw-ipd/RoseTTAFold2NA and running the command “run_RF2NA.sh rna_pred protein.fa R: RNA.fa”.

The paired homologous sequences generated by RoseTTAFoldNA during the prediction phase were subsequently employed to compute the coevolution score using EV_RNA^32^, a tool obtained from https://github.com/debbiemarkslab/plmc. This score serves as an indicator of the likelihood of RNA-protein contacts. PractIP^31^, sourced from https://github.com/satoken/practip, was locally installed and applied for contact prediction based on sequence and predicted secondary structure data. The tool to predict RNA and protein secondary structure is installed following the requirements of PractIP. It yields a binary contact map directly. However, this format is not conducive to calculating the auROC during evaluation, as it does not provide a probability matrix. Consequently, PractIP’s output can only contribute a single data point on the ROC curve.

To produce ROC curves for AlphaFold 3 and RoseTTAFoldNA, the distances between nucleotides and residues in the first complex model predicted by AlphaFold 3 or RoseTTAFoldNA were extracted and transformed to the probability of the contacts [**eq. (8)**] as below, as has been done by ^35,62,63^:

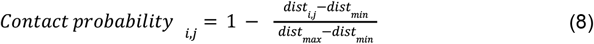

Here, *dist*_*i,j*_ represents the distance between the ith nucleotide and the jth residue, with *dist*_*min*_ and *dist*_*max*_ denoting the minimum and maximum distances, respectively, across all nucleotide-residue pairing combinations within a given predicted RNA-protein complex. The max-min normalization will not alter the ranking of the distances between nucleotides and residues.

## Code Availability

The open-source code of RPcontact is available at https://github.com/JulseJiang/RPcontact

## Data availability

All data are freely available from the Protein Data Bank (PDB) at https://www.rcsb.org/. The split of datasets, RNA-protein list, and the statistics of these datasets are provided in the Supplementary Data file (Supplementary_data.xlsx).

## Acknowledgement

The work at Shenzhen Bay Laboratory is supported in part by the computing facility at Shenzhen Bay Laboratory and Shenzhen Medical Academy of Research and Translation. Similarly, the work at the Guangzhou National Laboratory has benefited from the computational infrastructure available at the Guangzhou National Laboratory.

## Author Contributions

Y. Z. & Z. M. conceptualized, supervised the study, acquired funding and provided resources. J. J., X.Z., J. Z., & Z. M. contributed the methodology. J. J. conducted the investigation and curated data, J. J. performed formal analysis, visualization and provided software. J.J., Z.M. & Y.Z. wrote the original draft of the manuscript. All authors reviewed and edited the manuscript.

