## Supplementary File for "RPcontact: Improved prediction of RNA-protein contacts using RNA and protein language models"

* To whom correspondence should be addressed.

### Supplementary Figures

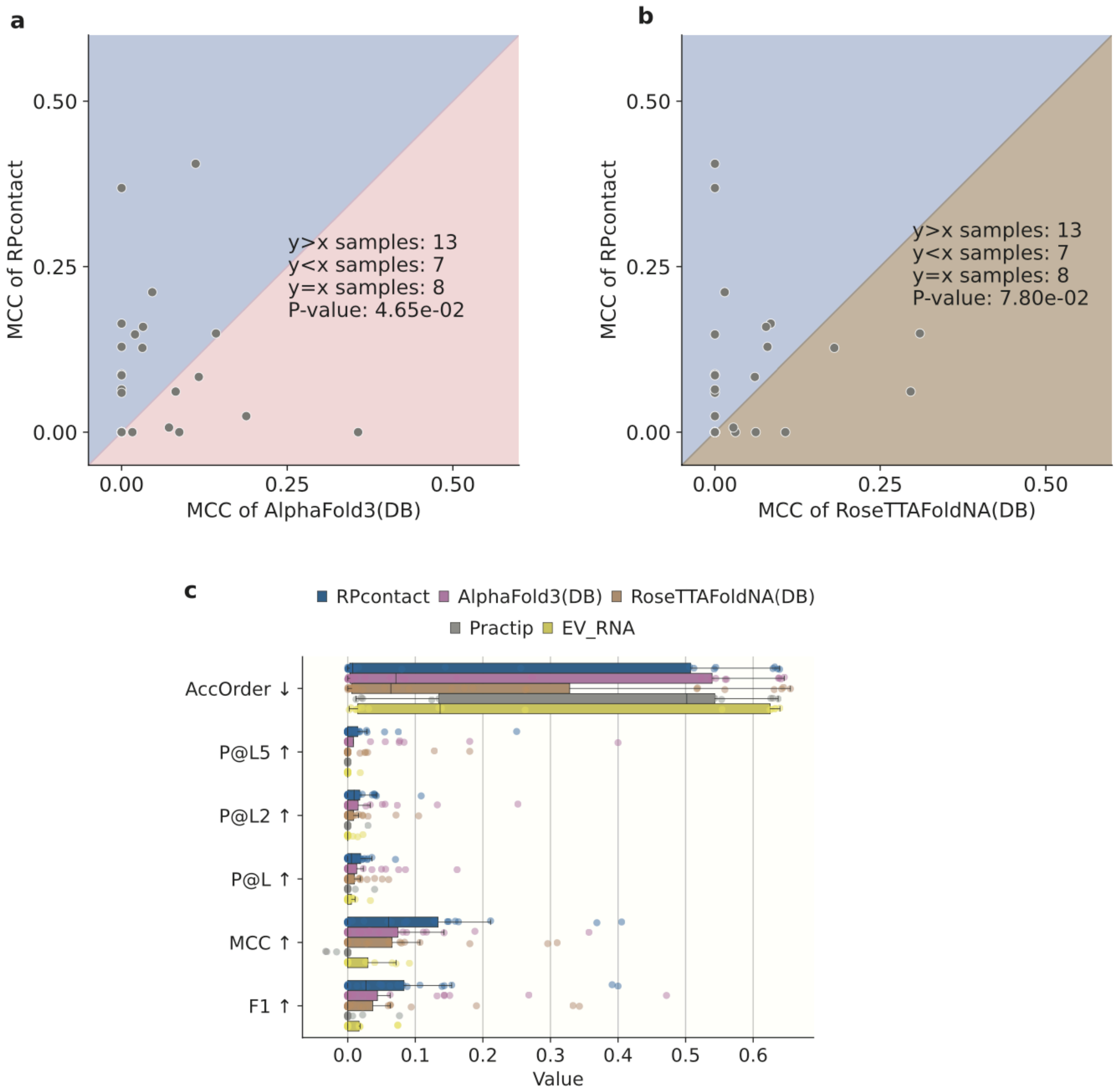


**Figure S1.** Robustness of RPcontact performance across multiple test sets, along with comparison to other methods, including AlphaFold 3(DB), RoseTTAFoldNA(DB), Practip, and EV_RNA on TS_nt.

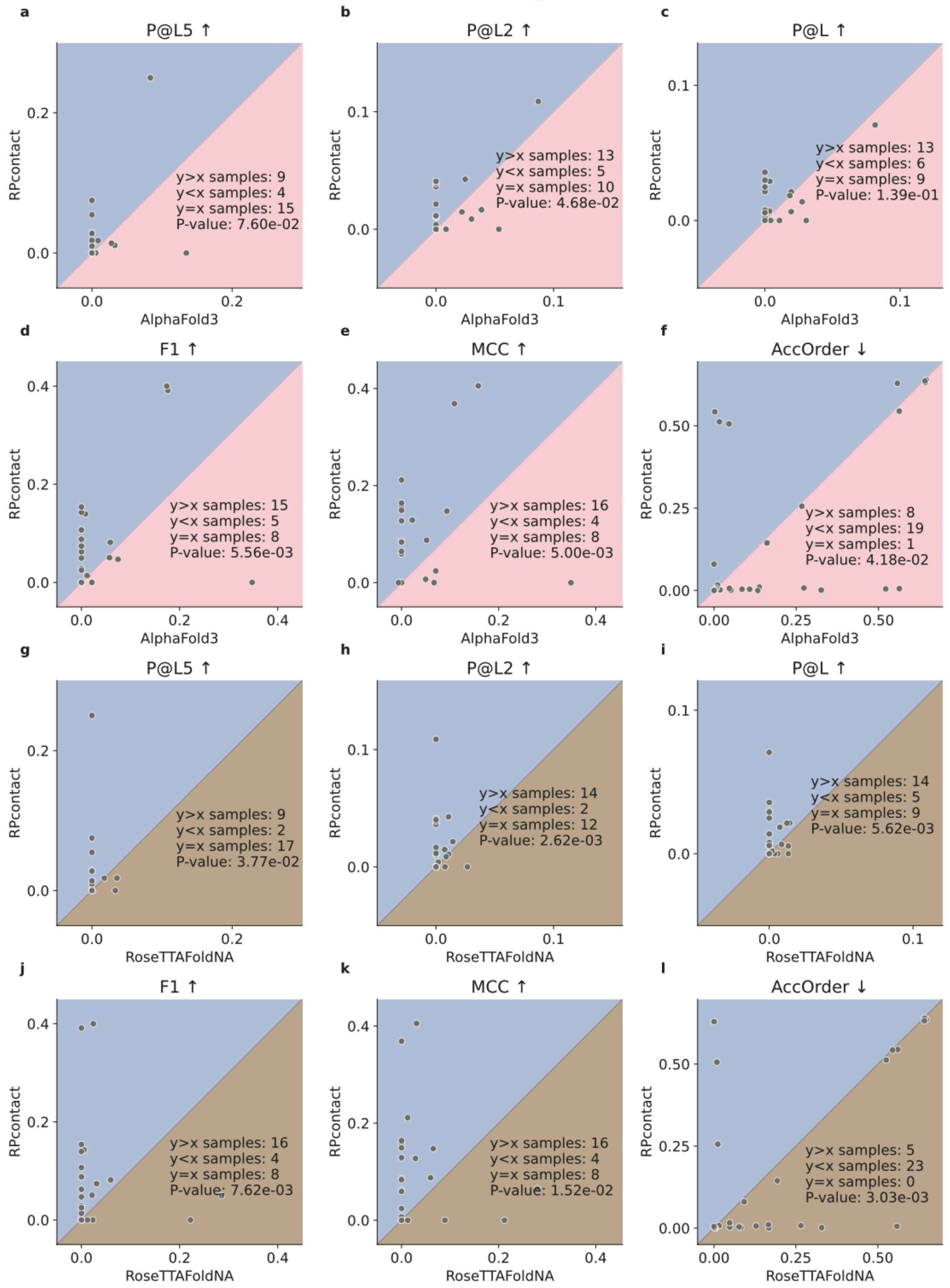


**Figure S2.** Comparing the performance of RPcontact with AlphaFold 3 and RoseTTAFoldNA utilizes diverse metrics evaluated on the TS_nt dataset.

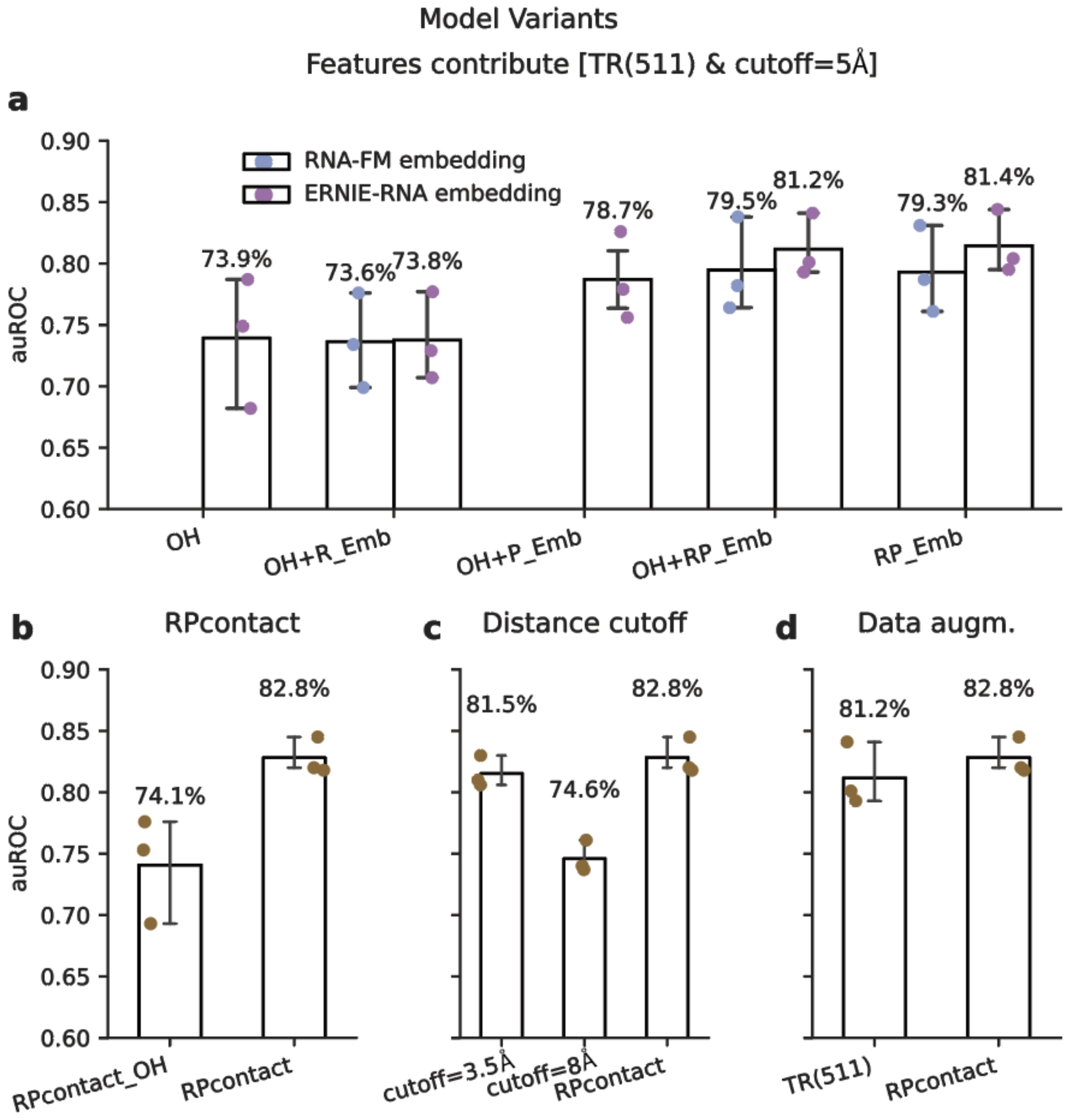


**Figure S3. Comparison of the auROC on a diverse dataset with different training settings. a.** Model performance with different embeddings (OH, OH+R_Emb, OH+P_Emb) and RNA embeddings (RNA-FM, ERNIE-RNA) at a 5 Å cutoff without data augmentation. **b.** Performance compared to RPcontact_OH with data augmentation but without LLM embeddings. **c**. Performance with labels generated by different distance cutoffs. **d**. Effect of data augmentation on model performance. The x-axis shows model variants, and the y-axis shows auROC evaluated at a 5 Å cutoff. Dots represent performance on VL, TS1, or TS2 datasets. The text above each rectangle indicates the average auROC across datasets, with error bars showing mean ± standard deviation. "+" means concatenating different features. LLM. Large language model.

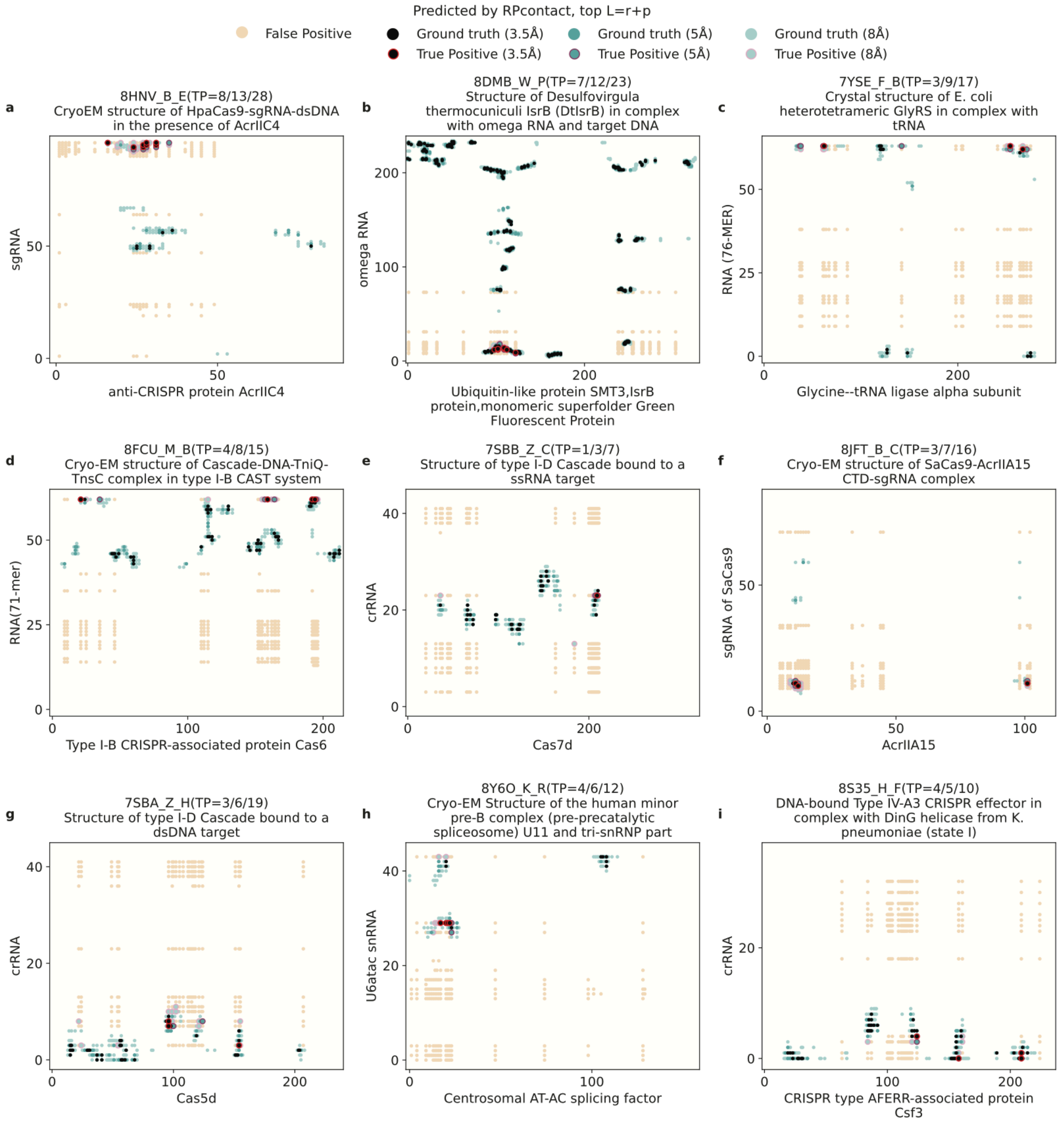


**Figure S4**. RPcontact's predictive accuracy for RNA-protein contacts across diverse complexes. Contact maps illustrate ground truth contacts, differentiated by green dot intensity corresponding to distance cutoffs of 3.5 Å, 5 Å, and 8 Å. The top L=r+p predicted contacts by RPcontact are also shown. True positive (TP) contacts are indicated with red to pink circles, varying with distance cutoffs, while yellow dots signify false positives. Each subplot corresponds to a distinct RNA-protein complex from the TS_nt dataset, with x- and y-axes denoting protein residues and RNA nucleotides. Complex names are derived from the PDB, and axis labels identify protein and RNA names. The notation TP=8/13/28 refers to the number of true positives for the respective distance cutoffs of 3.5 Å, 5 Å, and 8 Å.

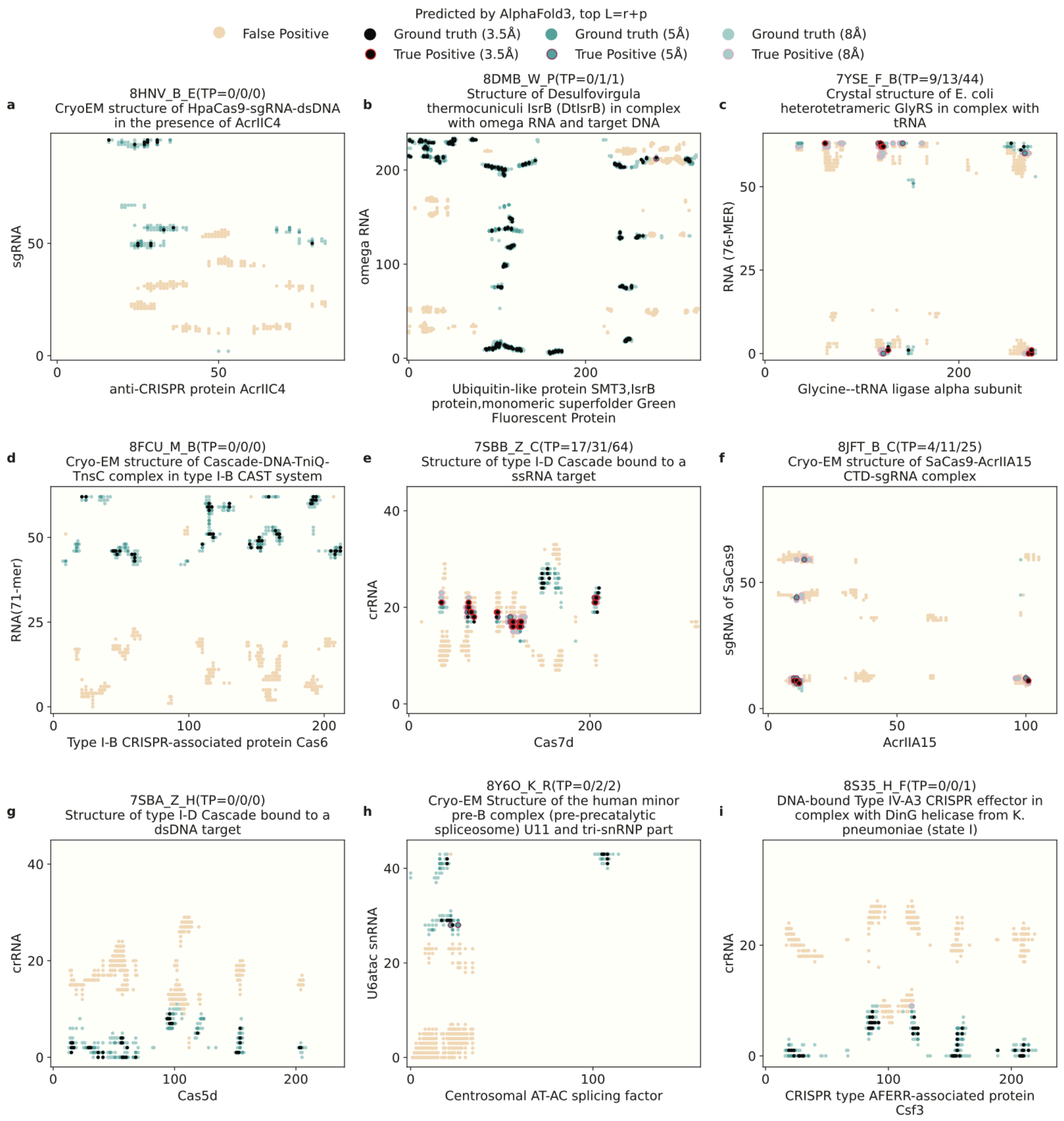


**Figure S5.** AlphaFold 3's predictive accuracy for RNA-protein contacts across diverse complexes.

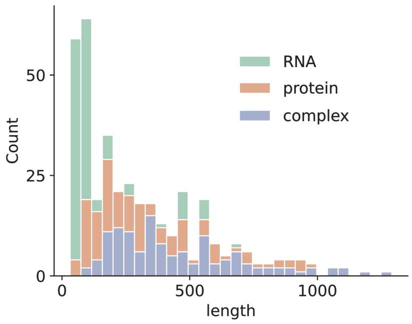


**Figure S6.** Sequence length distribution of RNAs, proteins, and complexes (the sum length of RNA and protein) in the combined test set (TS).

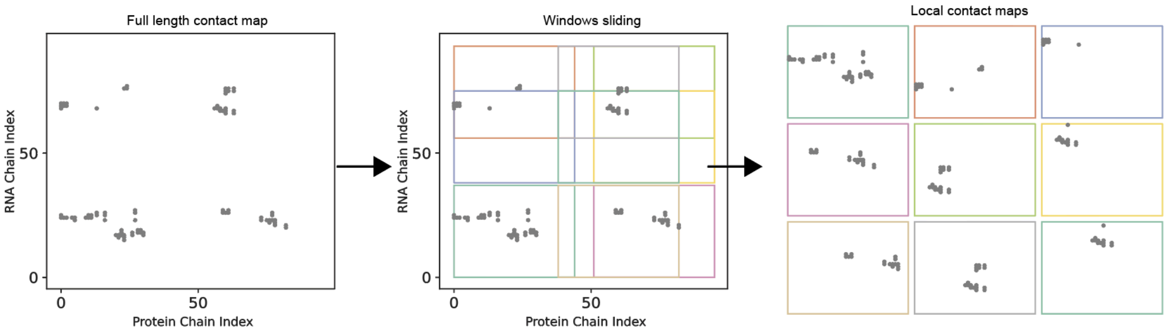


**Figure S7**. Illustration of the data augmentation in repeating local contacts for enhanced training in the regions with contacts.

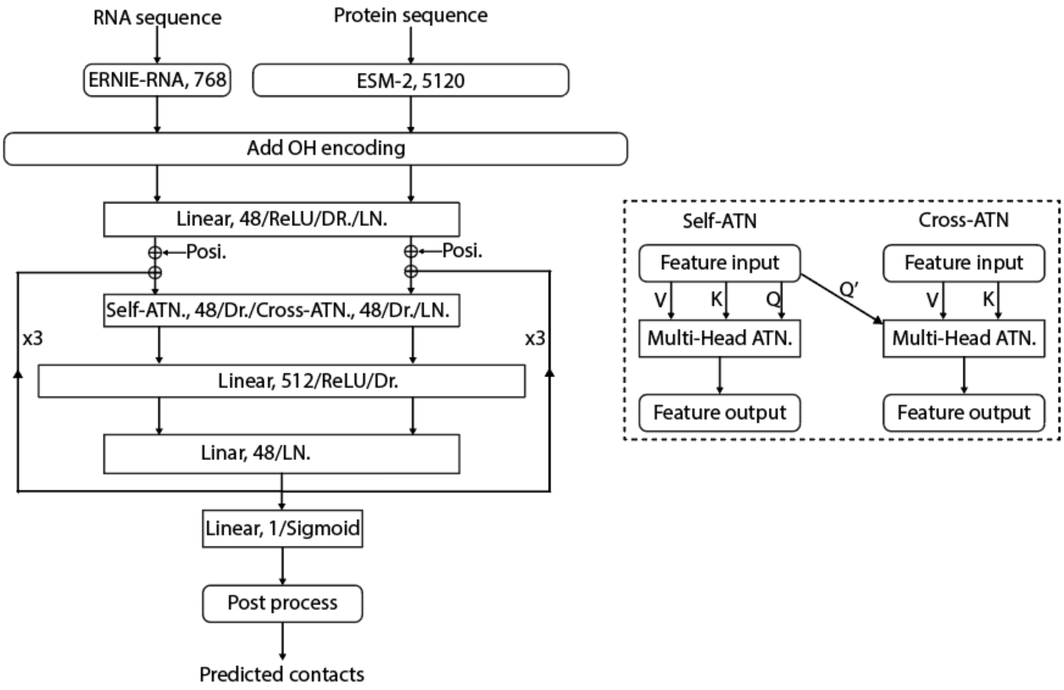


**Figure S8**. Details of the network structures in RPcontact.

### Supplementary Tables

**Table S1.** Statistics of the datasets (provided in Supplementary_data.xlsx)
